# LDJump: Estimating Variable Recombination Rates from Population Genetic Data

**DOI:** 10.1101/190876

**Authors:** Philipp Hermann, Angelika Heissl, Irene Tiemann-Boege, Andreas Futschik

**Affiliations:** Department of Applied Statistics, Johannes Kepler University Linz, Austria; Institute of Biophysics, Johannes Kepler University Linz, Austria

**Keywords:** Bioinformatics, population recombination rate, regression, change-point estimation, R-package

## Abstract

As recombination plays an important role in evolution, its estimation, as well as, the identification of hotspot positions is of considerable interest. We propose a novel approach for estimating historical recombination along a chromosome that involves a sequential multiscale change point estimator. Our method also permits to take demography into account. It uses a composite likelihood estimate and other summary statistics within a regression model fitted on suitable scenarios. Our proposed method is accurate, computationally fast, and provides a parsimonious solution by ensuring a type I error control against too many changes in the recombination rate. An application to human genome data suggests a good congruence between our estimated and experimentally identified hotspots. Our method is implemented in the `R`-package *LDJump*, which is freely available from https://github.com/PhHermann/LDJump.

## 1 Introduction

Recombination is a process during meiosis starting with the formation of DNA double-strand breaks (DSBs) and resulting in an exchange of genetic material between homologous chromosomes [Baudat et al., 2013]. The process leads to the formation of new haplotypes and increases the genetic variability in populations. In most species, recombination is concentrated in narrow regions known as hotspots, 1-2 kb in length, flanked by large zones with low recombination or cold regions. Meiotic recombination is a tightly regulated process defined mostly by a methyltransferase protein called PR domain zinc finger protein 9 (PRDM9) in most mammals (reviewed in [Baudat et al., 2013, Tiemann-Boege et al., 2017]). PRDM9 binds to a certain sequence motif (Myers motif) with its zinc finger array and recruits the DSB machinery (SPO11) to the hotspot (reviewed in [Tiemann-Boege et al., 2017]). Hotspots vary between species (human vs. chimpanzee see [Auton et al., 2012] or mice see [Smagulova et al., 2011]), populations within species (human populations like Africans and Europeans see [The 1000 Genomes Project Consortium, 2015, Pratto et al., 2014, Berg et al., 2010]), individuals within species (humans see [Pratto et al., 2014]), individuals of different sexes (see [Kong et al., 2010]) as well as between viruses (reviewed in [Pérez-Losada et al., 2015]).

Molecular and evolutionary mechanisms of the process of recombination can be better understood with accurate local estimates of the recombination rate [McVean et al., 2004, Chan et al., 2012]. Moreover, knowledge of the recombination rate variation along DNA sequences improves inference from polymorphism data about e.g. positive selection [Sabeti et al., 2006], linkage disequilibrium [Hill and Robertson, 1968], and facilitates an efficient design and analysis of disease association studies [McVean et al., 2004]. For this purpose, we designed *LDJump*, an algorithm that provides a fast and reliable new estimate of variable genome-wide historical recombination rates by partitioning the DNA sequence into homologuous regions with respect to recombination that also permits to take demography into account.

Methods differing in their genome-wide coverage, resolution, and reliance on active recombination have been proposed to estimate recombination rates in humans. Experimental approaches include whole genome sequencing or SNP typing of pedigrees of at least 2-3 generations [Coop et al., 2008, Kong et al., 2010, Halldorsson et al., 2016], leading to a resolution of order tens of kilobases, given that not many recombination events are captured. Direct measurements in sperm provide high resolution events at the level of a few hundreds of base pairs, but lack genome-wide coverage [Kauppi et al., 2004, Arnheim et al., 2007, Arbeithuber et al., 2015]. Finally, recombination hotspots have been inferred by the analysis of patterns of linkage disequilibrium [McVean et al., 2004, Myers et al., 2005, Myers et al., 2008]. The latter approach provides genome-wide historical recombination events based on polymorphisms characterized in several individuals within a population.

One of the first approaches to infer the population recombination rate *ρ* from patterns of linkage disequilibrium was to compute a lower bound on the number of recombination events [Hudson and Kaplan, 1985, Wiuf, 2002, Myers and Griffiths, 2003]. In population genetics, *ρ* is defined as *ρ* = 4*N_e_r*, where *N_e_* is the effective population size and *r* the recombination rate per base pair (bp) and generation. Other methods estimate *ρ* via maximum likelihood [Kuhner et al., 2000, Fearnhead and Donnelly, 2001] or approximations to the likelihood [Hudson, 2001, Fearnhead and Donnelly, 2002, McVean et al., 2002, Li and Stephens, 2003, Wall, 2004]. The former methods rely on simulations using importance sampling [Fearnhead and Donnelly, 2001] or Markov chain Monte Carlo (MCMC) methods [Kuhner et al., 2000] to become computationally feasible. The latter approaches use a composite likelihood as in [Hudson, 2001], or a modified composite likelihood as in [McVean et al., 2002]. Software implementations such as *LDhat* [McVean et al., 2004, Auton and McVean, 2007] and *LDhelmet* [Chan et al., 2012] are also available. [Kamm et al., 2016] extend this approach to account for demographic effects in their software package *LDpop*. Generally, computing approximate likelihoods requires a somewhat smaller computational effort than full likelihoods at the price of a slight loss in accuracy. An improvement of composite likelihood estimators via optimizing the trade-off between bias and variance has been proposed by [Gärtner and Futschik, 2016]. For a more technical discussion on composite likelihood in general see [Varin et al., 2011, Reid, 2013].

Another approach is to rely on calculating moments or summary statistics [Hudson, 1987, Batorsky et al., 2011]. In [Wall, 2000, Wall, 2004], suitably chosen summary statistics such as the number of haplotypes (*haps*) are used. There the author performs simulations with given *haps*, calculates the likelihood for a series of *ρ* values, and chooses the value of *ρ* with the highest likelihood as estimator of the recombination rate.

Further well-established frameworks to estimate recombination rates include `Lamarc` [Kuhner, 2006], `OmegaMap` [Wilson and McVean, 2006], `RDP` [Martin et al., 2015], and `CodABC` [Arenas et al., 2015]. The latter method [Arenas et al., 2015] applies ABC (approximate Bayesian computation) using 26 summary statistics to estimate constant recombination rates for simulated regions of size up to 300 codons for 100 alignments. With the GUI of `RDP` [Martin et al., 2015] overall patterns of recombination and testing for hot- and coldspots is performed by integration of *LDhat* [McVean et al., 2004]. Recently, alternative fast estimates of *ρ* that rely on regression on sliding windows have been proposed by [Lin et al., 2013, Gao et al., 2016]. Their software implementation is called *FastEPRR* and recommended for larger samples consisting of 50 sequences or more.

So far all these previous methods have at least some limitations such as being computationally demanding, not designed for small sample sizes or leading to a too large number of breakpoints in the recombination map. In order to address these issues within one algorithm we developed *LDJump*. More precisely, we divide the DNA sequence into small segments and estimate the recombination rate per segment via a regression based on the following summary statistics: measures of LD, *r*^2^, Watterson’s *θ*, measures on pairwise differences, haplotype heterozygosity, the four gametes test as well as the constant recombination rate estimator of *LDhat* [McVean et al., 2004]. A frequentist segmentation algorithm [Frick et al., 2014] is then applied to the estimated rates to obtain change-points in recombination. The algorithm controls a type I error and provides confidence bands for the estimator. [Futschik et al., 2014] use a similar approach to partition DNA sequences into homogeneous segments with respect to GC content. In contrast to [Gao et al., 2016] our approach is also designed to work with small sample sizes. As will be shown in the following sections, *LDJump* allows us to calculate hotspots at high accuracy within a reduced computational time from sample sizes of at least 10 sequences of genomic regions spanning many megabases.

Section 2 contains a detailed description of our proposed method we call *LDJump*. In section 3 we assess *LDJump* and compare it with the popular software packages *LDhat, LDhelmet*, and *FastEPRR*. We also consider different levels of genetic diversity as well as demographic effects. For different human populations, we apply our approach to a well-characterized region of the human genome. We furthermore estimate population specific recombination maps for the complete human chromosome 16, showing a good overlap between our and experimental estimates of hotspot positions. Finally, we summarize our findings in section 4. Further details on the regression model, bias correction, and more detailed simulation results are provided in the supplementary material.

## 2 Materials and Methods

Our approach consists of two steps. First, we fit a regression model from simulated data to estimate constant recombination rates on small segments. Subsequently, we apply a segmentation algorithm to estimate breakpoints in the recombination rate which is subject to type I error control against over-estimating the number of identified breakpoints.

### 2.1 Regression Model for Constant Recombination Rates

We use generalized additive models (GAM) [Wood, 2011] to estimate cubic spline functions *f_j_*(*Z_j_*) for covariates *z_j_, j* = 1,…, *q* and linear (or quadratic) effects for covariates *x_k_,k* = 1,…, *l* to regress the population recombination rate *ρ* on summary statistics computed from simulated short DNA segments. The structure of our GAM is

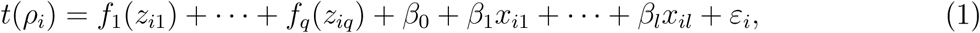
for *i* = 1,…, *n*. We chose our explanatory variables using an ANOVA on simulated data. The resulting set of (suitably scaled) summary statistics *χ* includes the constant recombination rate estimator available within the *LDhat* package, and can be found in Table 1.

**Table 1:**
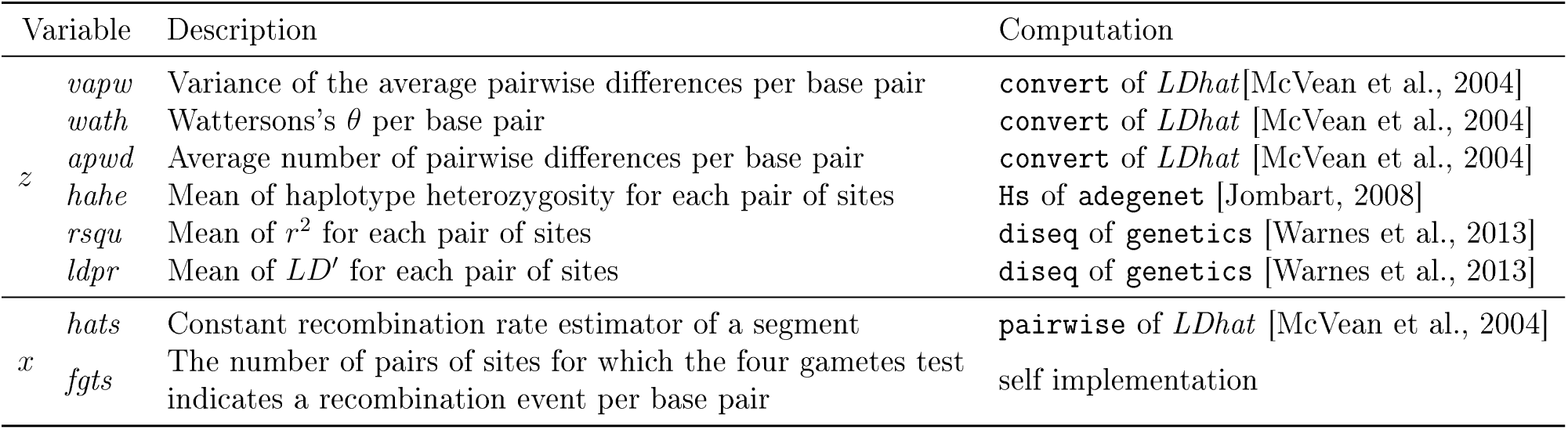
Summary statistics used in the regression model. The influence of variables tagged with *z* is modeled using spline functions and variables tagged with *x* are fitted by linear and quadratic effects.

For a more detailed description of the regression model as well as the selection of explanatory variables see supplementary material section 1.1. Initial computations revealed variance heterogeneity. Hence, we transformed the population recombination rate *ρ* using a Box-Cox transformation *t*(*ρ*) [Box and Cox, 1964]. In section 1.2 of the supplementary material, we describe the choice of the transformation parameters.

We observed a systematic overestimation of the background rates as well as underestimated hotspot intensities. Therefore, we performed a simulation based bias correction using quantile regression of the true recombination rate on the above described estimates. For further details on the bias correction see Figure 3 and section 1.3 of the supplementary material.

### 2.2 Segmentation Algorithm Estimating Variable Recombination Rates

[Frick et al., 2014] introduced a method called SMUCE for detecting change points in a function for observations distributed according to an exponential family. This method starts with a constant function and introduces successively additional jumps, as long as they lead to a significant increase in the likelihood. Using likelihood ratio tests, the probability of overestimating the number of change-points is controlled subject to a user specified type I error probability *α*. For a given number of jumps, the best fitting locally constant function is chosen by maximizing the likelihood. We use this method with local estimates 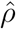 as input. For a general overview on multiple change-point detection see [Niu et al., 2016].

In the first step *LDJump* divides the DNA sequence into *k* short segments. Summary statistics are computed separately for each segment and inserted into our regression model to estimate a local transformed recombination rate. The back-transformed rates follow an approximate normal distribution (natural scale of *ρ*, see supplementary material section 1.2) and are used as input for the change point estimator. In our simulations, the use of the back-transformed rates led to a better detection of hotspots compared to the transformed rates.

## 3 Results

We used the software package *scrm* of [Staab et al., 2014] to simulate samples of populations with variable recombination rates and converted its output to *fasta-*files with the software package *ms2dna* of [Haubold and Pfaffelhuber, 2013]. In this section we compare *LDJump* with *LDhat*, the newer version *LDhat2, LDhelmet* as well as *FastEPRR*. We consider both constant and variable recombination rates and look at the performance as well as the runtime. The runtime comparison is based on one core of an Intel Xeon E5-2630v3 2.4 1866, with 64GB DDR4-2133 RAM. Our analysis was performed in [R Development Core Team, 2017]. Note that all mentioned software packages can also be applied on several cores in parallel.

### 3.1 Constant Recombination Rate Estimation

We first focus on a constant recombination rate on a DNA segment. In our simulations, *LDJump* is compared with the functions `pairwise` of *LDhat* and `max_lk` of *LDhelmet* following the default guidelines. The chosen sample sizes were {10, 16, 20}, and the sequence lengths {1000, 2000, 3000} base pairs. For each of these nine setups we simulated under 111 different values of *ρ* ∈ [0, 0.1] yielding a total of 999 simulated scenarios. The population mutation rate was chosen *θ* = 0.01.

Using the root mean squared error 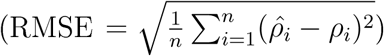 and the coefficient of determination *R*^2^, we compare the accuracy of the mentioned methods. We visualize the estimators and the true values in Figure 1 along with a diagonal black line indicating the true values. Both prediction measures show a slightly better fit of the generalized additive model (purple plus signs: higher *R*^2^ of 0.4974; smaller RMSE of 0.0256) compared with the software packages *LD-hat* (red dots: 0.4447; 0.0290) and *LDhelmet* (green triangles: 0.2095; 0.0360). As our method uses the function *pairwise* as one of the summary statistics, the improved performance may be in part due to an optimized bias-variance trade-off, see [Gärtner and Futschik, 2016].

**Figure 1:**
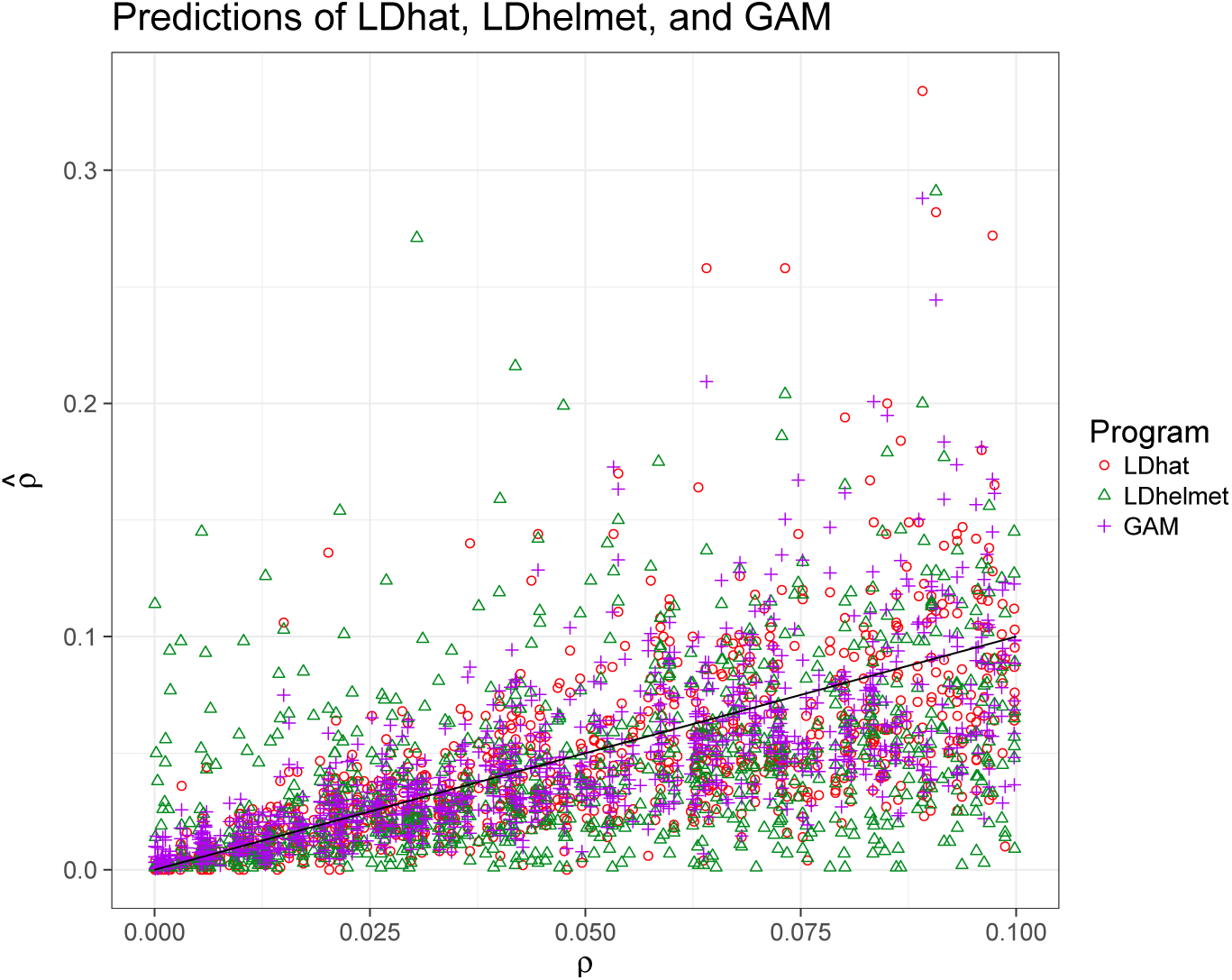
Predicted recombination rates versus their true values estimated with *LDhat* (red dots). *LDhelmet* (green triangles) and the GAM (purple plus signs). The black line provides the true rates.

### 3.2 Variable Recombination Rate Estimation

For humans, large fractions of recombination events are concentrated on short segments which are called hotspots (reviewed in [Arnheim et al., 2007]). Following the literature, we define recombination hotspots as genomic regions that exceed the background rate by more than a threshold factor of five for a length of up to 2kb [McVean et al., 2004].

We investigate how well hotspots are detected by our method and simulated two types of setup for variable recombination rate estimation: *simple* setups (sequences of length 10 and 20 kb with one hotspot) and *natural* setups (sequences of length 1Mb containing 15 hotspots) both using a mutation rate *θ* of 0.01. These scenarios were investigated with different background rates, sample sizes, hotspot intensities, and hotspot lengths. When comparing our approach with *LDhat*(*2*) and *LDhelmet*, we followed recommendations and used 10^6^ iterations for the reversible-jump MCMC procedure, sampled every 4000 iterations, chose a burn-in of 10^5^, and different block penalties of 0, 5, and 50. For the computations with *LDhelmet*, we used a window size of 50 SNPs, and 11 Padè coefficients. Results for *FastEPRR* were obtained using *winLength*=*stepLength* (segment lengths) of 500, 1000, 1500, and 2000 base pairs. We applied the implemented function `smuceR` within the R-epackage `stepR` [Hotz and Sieling, 2016] to estimate the change-points for our method.

#### 3.2.1 Simple Setups

We simulated samples of sizes {10, 16, 20} with sequence lengths of 10 kb and 20 kb. Our 15 considered background recombination rates were chosen equidistantly within [0.001, 0.03].

We considered hotspot intensities of {5, 10, 15, 20, 40}-fold the background recombination rate. The length of the hotspots varied among 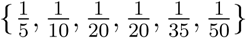-times the sequence length. Due to the large number of resulting setups and the computation times of *LDhelmet* and *LDhat(2)*, we have restricted this analysis to 2 replicates per sample yielding in total 4500 simulated recombination maps. We approximated the RMSE (root mean squared error) as our quality measure, and computed the estimation errors on an equidistant grid of 1000 positions along the sequences.

Table 2 summarizes the performance of the considered methods. More specifically, we computed the mean, median, and standard deviation (across simulations) of the RMSE for *LDhat* (column 3), *LDhat2* (c. 4), *LDhelmet* (c. 5), *FastEPRR* (c. 6-9, with different segment lengths) and *LDJump* (c. 10-15 with different numbers of user-defined segments *k*). The results using different block penalties for *LDhat*(*2*)*, LDhelmet* along with different type I error probabilities for *LDJump* are listed in separate rows.

**Table 2:** Mean 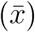, median (*x*_0.5_) and SD of the RMSE for *LDhat, LDhat2, LDhelmet* (LDhel), *FastEPRR*, and *LDJump* under *simple* setups. Different block penalties (*bpen*) have been tried for *LDhat, LDhat2, LDhelmet*. Different segment lengths have been applied with *FastEPRR*, and different number of segments as well as type I error probabilities *α* considered for *LDJump*.

| | *bpen* | *LDhat* | *LDhat2* | *LDhel* | *FastEPRR* (Segment Length) | | | | $\alpha$ | *LDJump* (Number of Segments) | | | | |
| --- | --- | --- | --- | --- | --- | --- | --- | --- | --- | --- | --- | --- | --- | --- |
|  |  |  |  |  | 500 | 1000 | 1500 | 2000 |  | 10 | 15 | 20 | 25 | 30 |
| $\bar{x}$ | 0 | 0.158 | 0.064 | 0.286 | | | | | 0.1 | 0.068 | 0.065 | 0.062 | 0.064 | 0.066 |
|  | 5 | 0.132 | 0.064 | 0.234 | 0.053 | 0.057 | 0.061 | 0.063 | 0.05 | 0.068 | 0.065 | 0.062 | 0.064 | 0.066 |
|  | 50 | 0.078 | 0.064 | 0.094 |  |  |  |  | 0.01 | 0.068 | 0.065 | 0.062 | 0.064 | 0.066 |
| $x_{0.5}$ | 0 | 0.138 | 0.036 | 0.247 | | | | | 0.1 | 0.042 | 0.040 | 0.036 | 0.037 | 0.040 |
|  | 5 | 0.100 | 0.036 | 0.169 | 0.039 | 0.034 | 0.035 | 0.036 | 0.05 | 0.042 | 0.040 | 0.036 | 0.037 | 0.040 |
|  | 50 | 0.049 | 0.036 | 0.044 |  |  |  |  | 0.01 | 0.042 | 0.040 | 0.036 | 0.037 | 0.040 |
| SD | 0 | 0.115 | 0.076 | 0.227 |  |  |  |  | 0.1 | 0.077 | 0.074 | 0.096 | 0.078 | 0.079 |
|  | 5 | 0.121 | 0.076 | 0.224 | 0.053 | 0.066 | 0.072 | 0.074 | 0.05 | 0.077 | 0.074 | 0.096 | 0.078 | 0.079 |
|  | 50 | 0.102 | 0.076 | 0.145 |  |  |  |  | 0.01 | 0.077 | 0.075 | 0.096 | 0.078 | 0.079 |

As discussed in the supplementary material (section 1.4), segment lengths of at least 333 bp are needed with *θ* = 0.01 for a good performance of *LDJump*. Following this recommendation, our method performs equivalently or slightly better than *LDhat2*, and outperforms *LDhat* and also *LDhelmet*. The choice of *α* did not have a large effect under the considered scenarios. Similarly, the block penalty does not much affect the performance of *LDhat2*. With *LDhat* and *LDhelmet* on the other hand, the choice of the block penalty strongly influences the performance. The performance of *LDJump* and *LDhat2* turned out to be more constant across simulations. Indeed, the standard deviation of the RMSE is more than 20 % lower with *LDJump* than that of *LDhat*, which in turn has a more than 30 % lower SD than *LDhelmet*. With FastEPRR, approximately 57%, 5%, 4%, and 2% of the computations terminated due to errors using segment lengths of 500, 1000, 1500, 2000, respectively. When *FastEPRR* provided estimates, the performance was comparable with *LDJump*. A more detailed graphical display of the performance of *FastEPRR* with respect to segment lengths can be found in Figure 7 in section 2 of the supplementary material.

Figure 2 contains separate results for different sample sizes, recombination rates, hotspot intensities and lengths, as well as sequence lengths. We applied *LDJump* with 20 segments and a type I error probability of 5%. Hence, the considered segments had a length of 500 and 1000 (for 10kb and 20kb, respectively) nucleotides. We used *FastEPRR* with a window length of 2kb in order to achieve a small number (32) of runs terminating due to errors. Especially for small to middle background rates (under the considered values) *LDJump, FastEPRR*, and *LDhat2* have on average a lower RMSE than *LDhat* and *LDhelmet. LDJump, FastEPRR* and *LDhat2* lead on average to a smaller RMSE for all considered sample sizes as well as sequence lengths, with our approach performing best in many cases. Moreover, slightly smaller or equivalent values for the RMSE were computed with *LDJump* and *FastEPRR* for hotspot intensities from 5- to 20-fold the background recombination rate and hotspot lengths between 1/50 and 1/10. For a hotspot length of 1/5 of the total sequence length we can see a similar fit for all methods. *LDJump* and *FastEPRR* have similar estimation quality with slight preference for our method.

**Figure 2:**
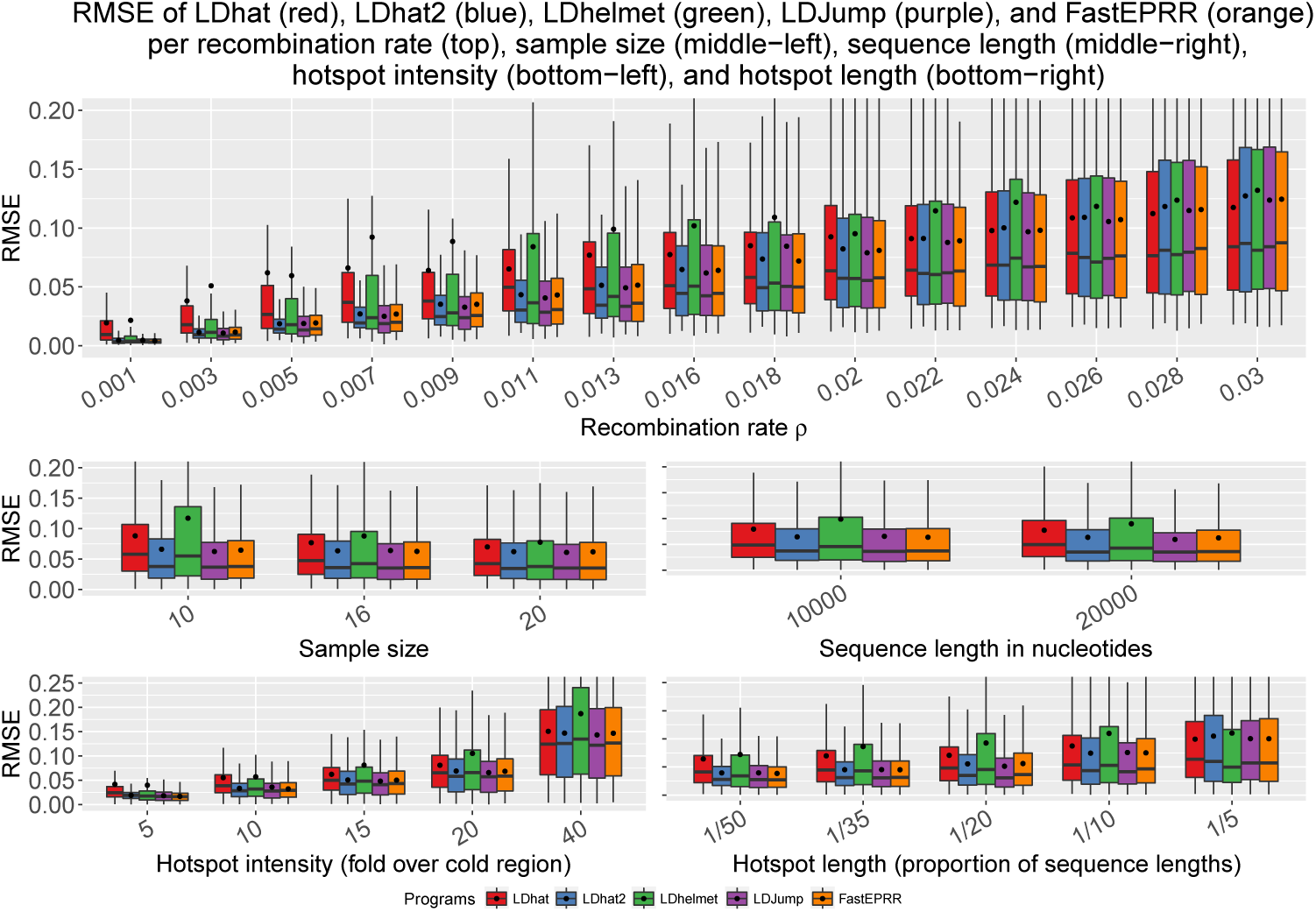
Comparison of the methods (*LDhat* (*red*)*, LDhat2*(blue). *LDhelmet* (green). *LDJump* (purple), and *FastEPRR*. (orange) for different true recombination rates (top), sample sizes (middle-left), sequence lengths (middle-right), hotspot intensities (bottom-left), and hotspot lengths (bottom-right). Mean values are shown as black dots.

#### 3.2.2 Natural Setups

We simulated samples with 16 sequences and sequence lengths of 1Mb. The setups varied in the background rate which was chosen among 13 equidistant values between 0.001 and 0.01. The 15 hotspots were evenly distributed along the sequence and had different intensities of 8 to 40-fold the background rate. Every setup was replicated 20 times. The same mutation rate *θ* = 0.01 was again chosen for all setups. In our simulations, we focused on the methods that performed best for the simple scenarios. As *FastEPRR* using segment lengths of 1kb terminated without producing estimates for 88% of our simulated complex data sets, we mainly restricted our attention to a comparison between *LDJump* and *LDhat2*. Additional information on the performance of *FastEPRR* based on the non-terminating runs only can be found in supplementary material section 3. Notice however that a high proportion of missing results may lead to a biased quality assessment, if the missing probability depends on features of the data set that affect the performance of the estimate.

Figure 3 provides a comparison between *LDJump* with (grey) and without (purple) bias correction, and *LDhat2* (blue). Three samples with different background recombination rates of 0.001 (left), 0.0054 (middle), and 0.01 (right) are presented in dotted black lines. Segment lengths were chosen to be 1kb with the quantile chosen 0.35 in the bias correction (see supplementary material section 1.3) and a type-I error probability of 0.05. The bias-correction decreases the bias in the background rates and increases the intensities of the estimated hotspots.

**Figure 3:**
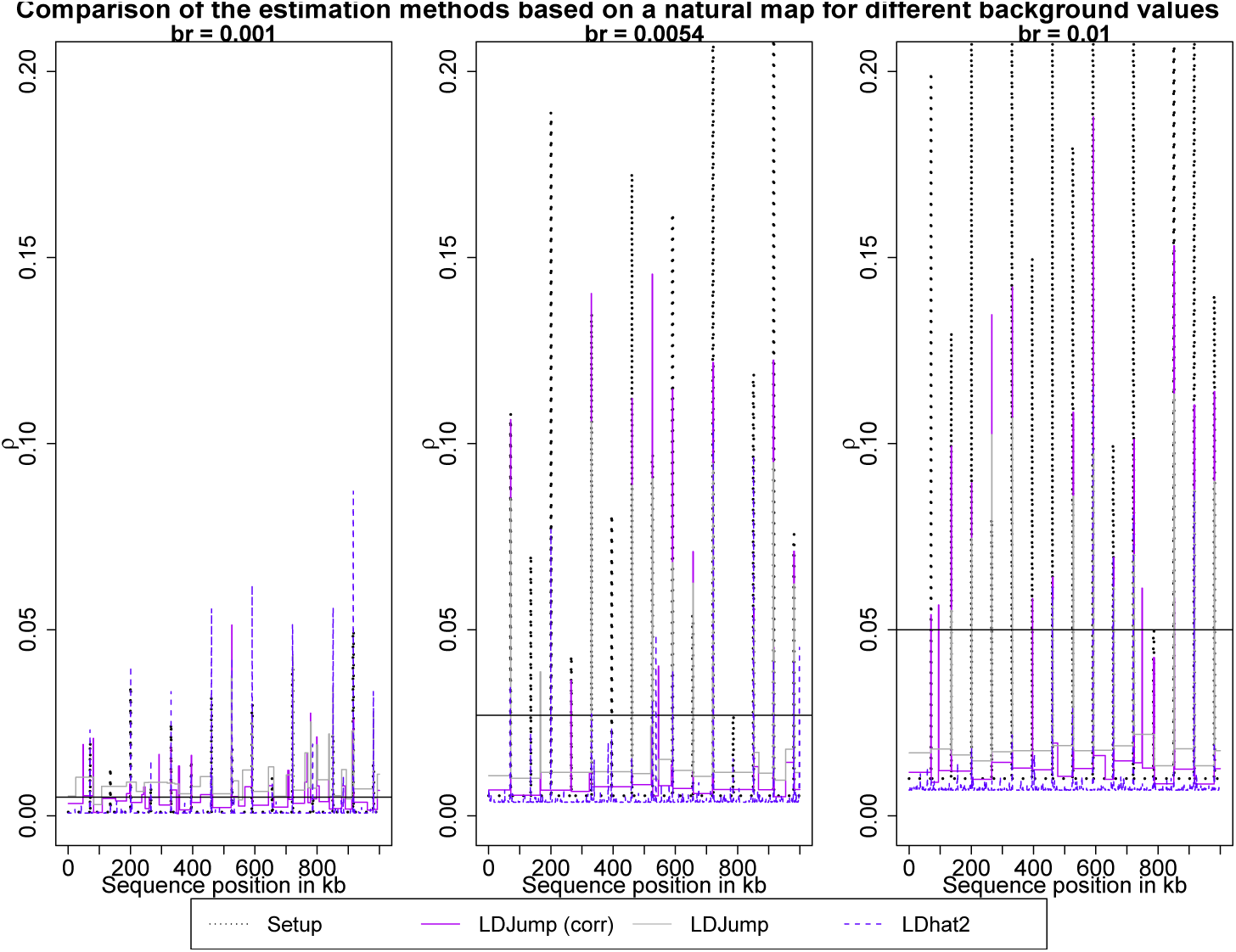
Estimated recombination maps using *LDhat'2* and *LDJump*, both with and without our bias-correction. The chosen setups differ in the background rates (0.001 (left). 0.0054 (middle), and 0.01 (right)). The true recombination map (black dotted lines) contains 15 hotspots. Horizontal lines represent the hotspot threshold (5-background rate).

**Quality Assessment** We took the weighted RMSE as measure of quality. It is defined as

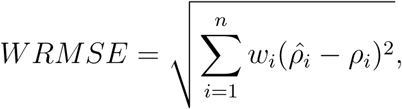
with *w_i_* denoting the length of the estimated segment *i* divided by the total sequence length. We also considered the proportion of correctly identified hotspots (PCH). A hotspot is counted as correctly identified if it has a non-empty intersection with a detected hotspot (i.e. a region with at least five-fold background recombination rate). The proportion of correctly identified background rates (PCB) has been defined analogously. Finally, the weighted average performance is given as WAP = (PCH + PCB)/2.

To identify the best combination of bias correction and segment lengths, we applied *LDJump* with *k* = 500, 1000, 1500, and 2000 segments and estimated the recombination maps using the 0.25, 0.35, 0.45, and 0.5 quantiles in the bias correction (see supplementary material section 1.3). Notice that segment lengths resulting from the chosen values of *k* are 2kb, 1kb, 666 and 500 bp. As hotspot lengths are either 1 or 2kb, the scenario with *k* = 1500 is most challenging as the hotspot boundaries will systematically differ from the segment boundaries. A direct comparison with *LDhat2* using a block penalty of 50 (based on the results from the *simple setups*) is provided.

The different choices of *k* are displayed by the first four groups of boxplots in Figure 4. For each of these four groups, quantiles of 0.25, 0.35, 0.4, and 0.5 are used in the bias correction and are presented in different colors. The rightmost bar in each panel (in blue) summarizes the result of *LDhat2*. From top-left to bottom-right, we show PCH, PCB, WAP, the estimated number of blocks, and the weighted RMSE.

**Figure 4:**
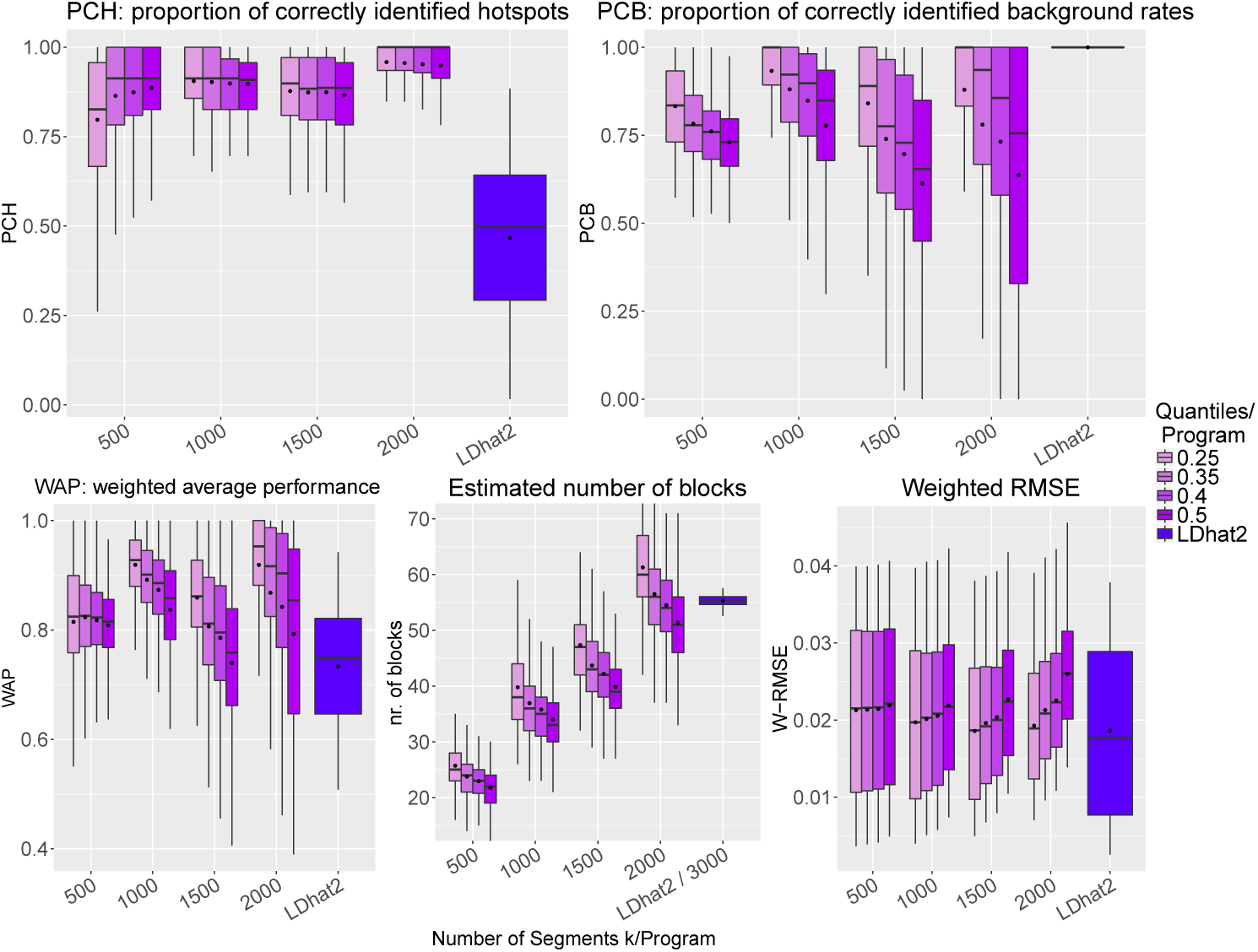
Quality assessment is performed based on the proportion of correctly identified hotspots (PCH, top-left), the proportion of correctly identified background rates (PCB, top-right), the weighted average performance (WAP = (PCH + PCB)/2, bottom-left), the estimated number of blocks (bottom-middle), and the weighted RMSE (bottom-right). Based on 13 setups with 20 replicates these measures are computed for *LDJump* using different numbers of initial segments *k* (500, 1000, 1500, 2000) and compared with the results of *LDhat2* using a block penalty of 50.

PCH may be interpreted as a measure of sensitivity, whereas PCB provides a measure of specificity. We can see that our method has very high detection rates irrespective of *k* with even less variability in performance than *LDhat2*. On the other hand, *LDhat2* has very high PCB proportions. The best PCB values for *LDJump* are obtained for the smallest quantile. As an overall measure, we display the mean of PCH and PCB as WAP in the bottom-left panel. It turns out that WAP is larger for *LDJump* regardless of the tuning parameters. In the bottom-middle panel we can see that the number of estimated blocks of *LDJump* depends on *k*. When using 500 segments, the estimated number of blocks is still below 31, the true number of blocks in the recombination map (due to 15 hotspots). For larger *k* the number of blocks is slightly overestimated. *LDhat2* estimated many more blocks, indeed the number of change points in recombination tended to be larger by a factor of more than 3000. The bottom-right plot shows the weighted RMSE as an overall quality measure showing a similar level of accuracy across *k* and compared with *LDhat2*. A more detailed investigation reveals that our method estimates hotspot rates more precisely, but provides less accurate estimators of the background recombination rate.

Our results also show that our method is fairly robust with respect to tuning choices. This is also true for *k* =1500, where the hotspots have an unfavorable location compared with the design segment boundaries. To obtain a reasonable tradeoff between sensitivity (PCH) and specificity (PCB), segment lengths of 1kb (based on 1000 segments of sequence length 1Mb) and a quantile of 0.35 in the bias correction seem to be a good choice with *LDJump*.

Figure 5 shows our considered quality measures depending on the background recombination rates. We provide the average performance over 20 replicates. We can see that *LDhat2* has constant PCB and decreasing PCH as the background rate increases. *LDJump* shows constant values for PCH and slightly increasing PCB for higher background rates. The overall measure WAP slightly increases for *LDJump* and decreases for *LDhat2* with increasing background rates, respectively. The weighted RMSE is also plotted. It can be seen that *LDhat2* leads to a slightly smaller weighted RMSE with decreasing differences for larger *ρ*.

**Figure 5:**
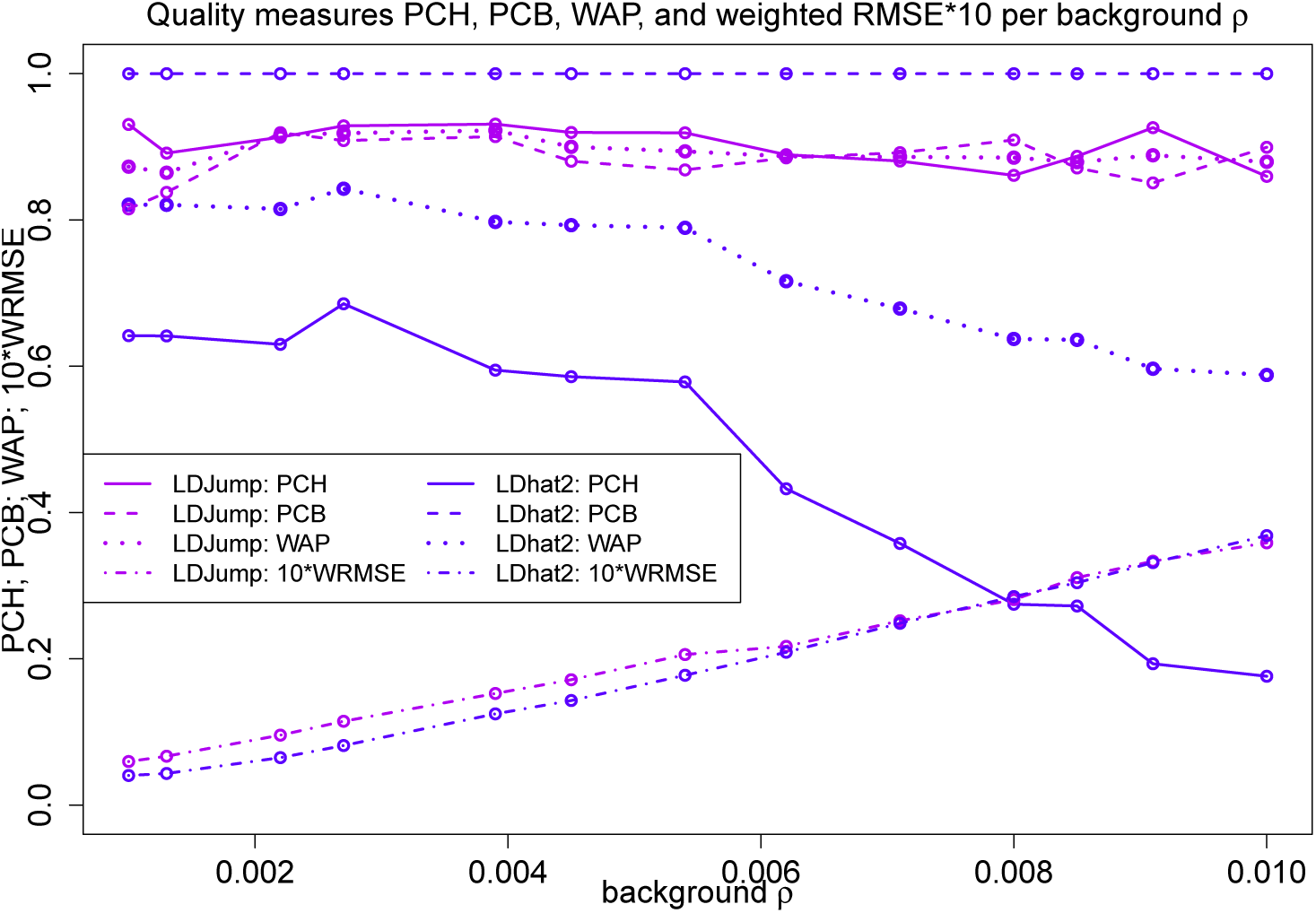
Proportion of correctly identified hotspots (PCH, solid), proportion of correctly identified background rates (PCB, dashed), the average of these two quality measures (WAP, dotted), and weighted RMSE*10 (dash-dotted) across different recombination rates. We compare *LDJump* (purple, segment length: 1kb, quantile 0.35). with *LDhat2* (blue, same line coding per quality measure).

Notice that we have obtained an error share of more than 88% using *FastEPRR* for the natural setups. We provide a comparison of the error-free results in Figure 8 in supplementary material section 3. Based on this smaller number of results for *FastEPRR*, *LDJump* is favorable in terms of the WRMSE and PCH, but has lower PCB compared to *FastEPRR*.

### 3.3 Populations under Different Levels of Genetic Diversity

Since natural populations differ in the level of genetic diversity, we simulated samples under different mutation rates *θ* ∈ = {0.0025, 0.005, 0.01, 0.02}. In Figure 6 we compare the performance based on the RMSE of *LDJump* (first panel) with *LDhat2*. For both methods, the influence of a misspecified *θ* has also been investigated. We used *LDJump* with segment lengths of 1kb, and the regression model calibrated under the mutation rate *θ* = 0.01. Thus the model is misspecified when the true *θ* ≠ 0.01. For *LDhat2*, results obtained using the true value of *θ* are displayed in the second panel, and results under misspecificationin the third panel.

**Figure 6:**
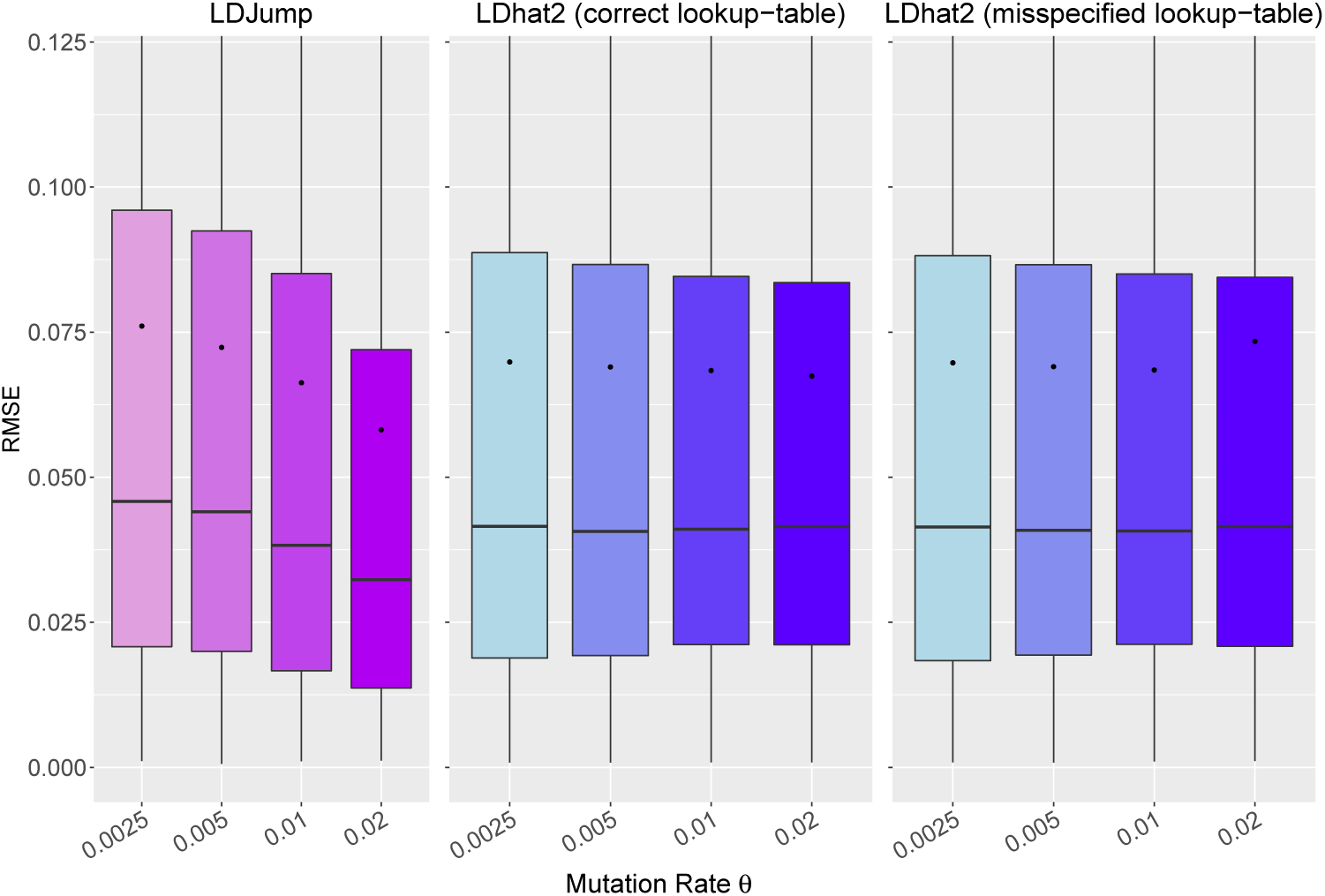
Accuracy of estimates for different levels of genetic diversity introduced by different mutation rates *θ*. *LDJump* (first panel) is compared with *LDhat2* for different values of *θ*. Misspecified values of *θ* are also considered: Indeed, *LDJump* was trained only under the mutation rate *θ* = 0.01. For *LDhat2*, we compare the performance under different mutation rates (2nd panel) and under misspecification assuming that the true mutation rate is equal to 0.01 (3rd panel, misspecified for *θ* ≠ 0.01).

*LDJump* improves with increasing mutation rates due to the higher information available per segment. Interestingly, *LDhat2* benefits less from increased levels of genetic diversity. A misspecified *θ* had little effect on the performance of *LDhat2*.

Based on our simulations we also evaluate the influence of the SNP density on the performance of *LDJump*. Figure 7 provides box plots illustrating the performance in terms of the RMSE depending on the mean number of SNPs per base pair within a simulated segment. Our results suggest that the higher the SNP density the more accurate estimates are obtained. When there are fewer than two SNPs in a segment, our software implementation imputes estimates based on the neighboring segments.

**Figure 7:**
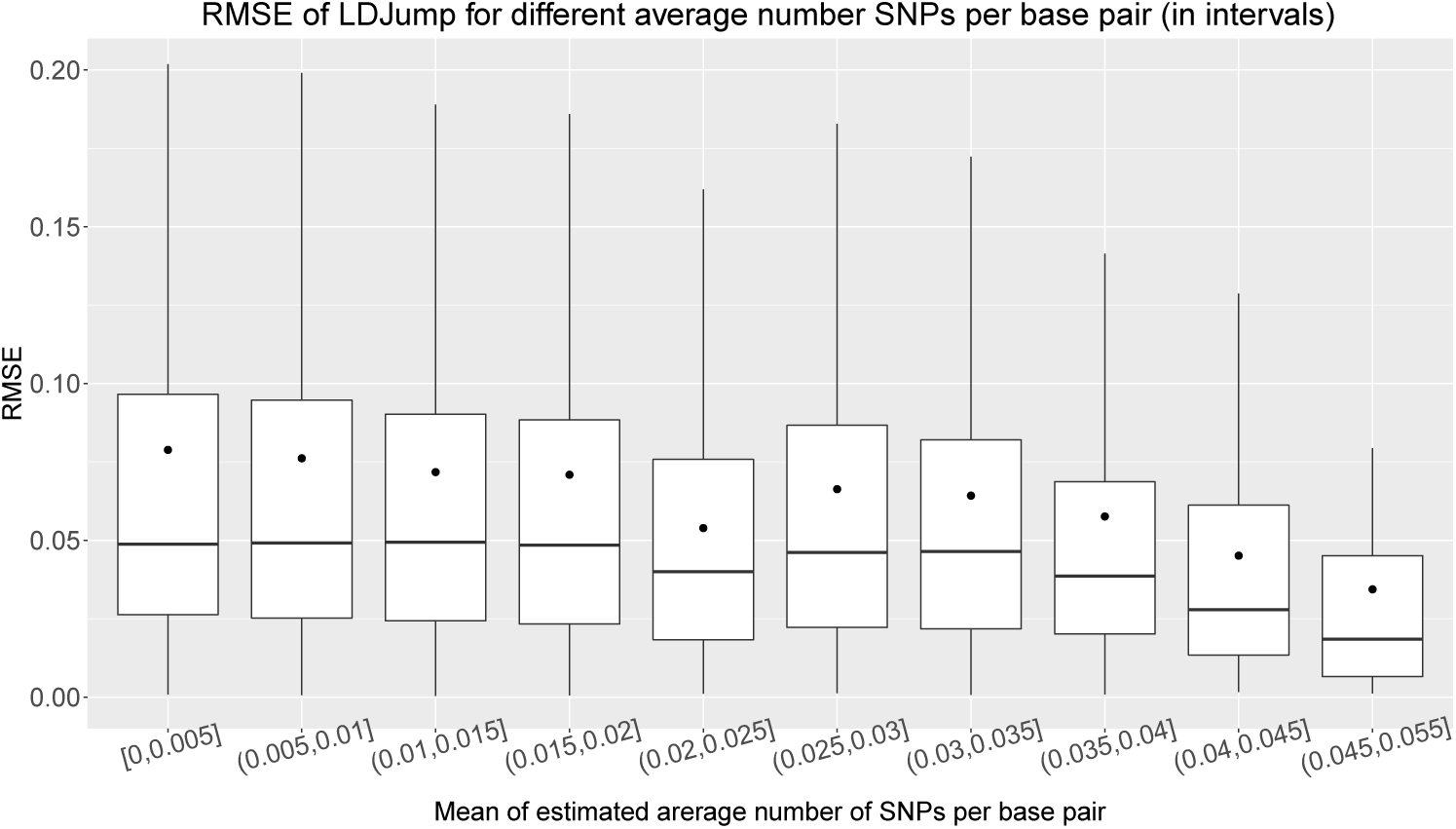
RMSE of *LDJump* using simulated populations under *simple* setups with different mutation rates *θ* = {0.0025, 0.005, 0.01, 0.02}. We compare the performance of *LDJump* in dependence on the mean number of SNPs per bp.

### 3.4 Populations under Demography

It has been observed in [McVean et al., 2002, Chan et al., 2012, Smith, 2005] that ignoring population demography by wrongly assuming a constant population size leads to biased estimates of recombination. As a remedy, [Kamm et al., 2016] computed 2-locus likelihoods under a known variable population size. *LDJump* permits the natural inclusion of any type of demography or even range of demographic scenarios by simply fitting our regression model under suitable scenarios.

We illustrate this approach and consider a scenario that involves a bottleneck followed by a rapid population growth. This scenario has also been used by [Kamm et al., 2016]. More precisely, we chose time dependent population sizes as follows:

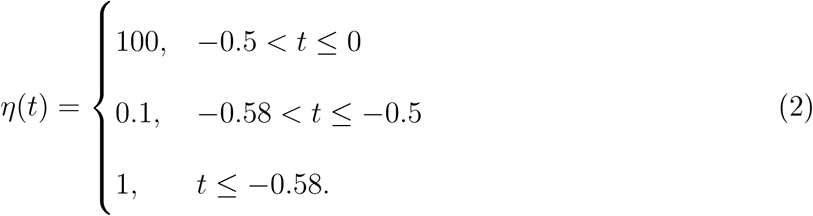

Time is scaled in coalescent units and the simulations were again performed with `scrm` [Staab et al., 2014]. [Johnston and Cutler, 2012] analyzed a similar demographic scenario and showed that *LDhat* infers spurious recombination hotspots when falsely assuming a constant population size.

With *LDJump*, we fitted our regression model using samples simulated under the demographic model (2). We used the same explanatory variables as under neutrality, but added Tajima’s D [Tajima, 1989] as an additional explanatory factor. This additional variable had significant effect on the model fit in our ANOVA, suggesting that choosing summary statistics dependent on demography can help to improve the accuracy of our estimates. We did not change the parameters in the Box-Cox transformation compared to the constant population size model. To see what can be gained by explicitly considering an underlying demography, we simulated samples under the demographic model (2), and sticking otherwise to the previously described *natural* setups. For these samples, we estimated recombination maps using the regression models trained either under neutrality (misspecified model) or under demography. More specifically *LDJump* has been applied with segment lengths of 1kb and a quantile of 0.35. The accuracy of these models was then compared in terms of the indicators PCH, PCB, and WRMSE.

The results are shown in Figure 8. When using the correct demographic model, the hotspot detection rate as well as the proportion of correctly identified regions with background recombination rate increases. We also found the WRMSE to be equal or slightly smaller when using the correct demographic model.

**Figure 8:**
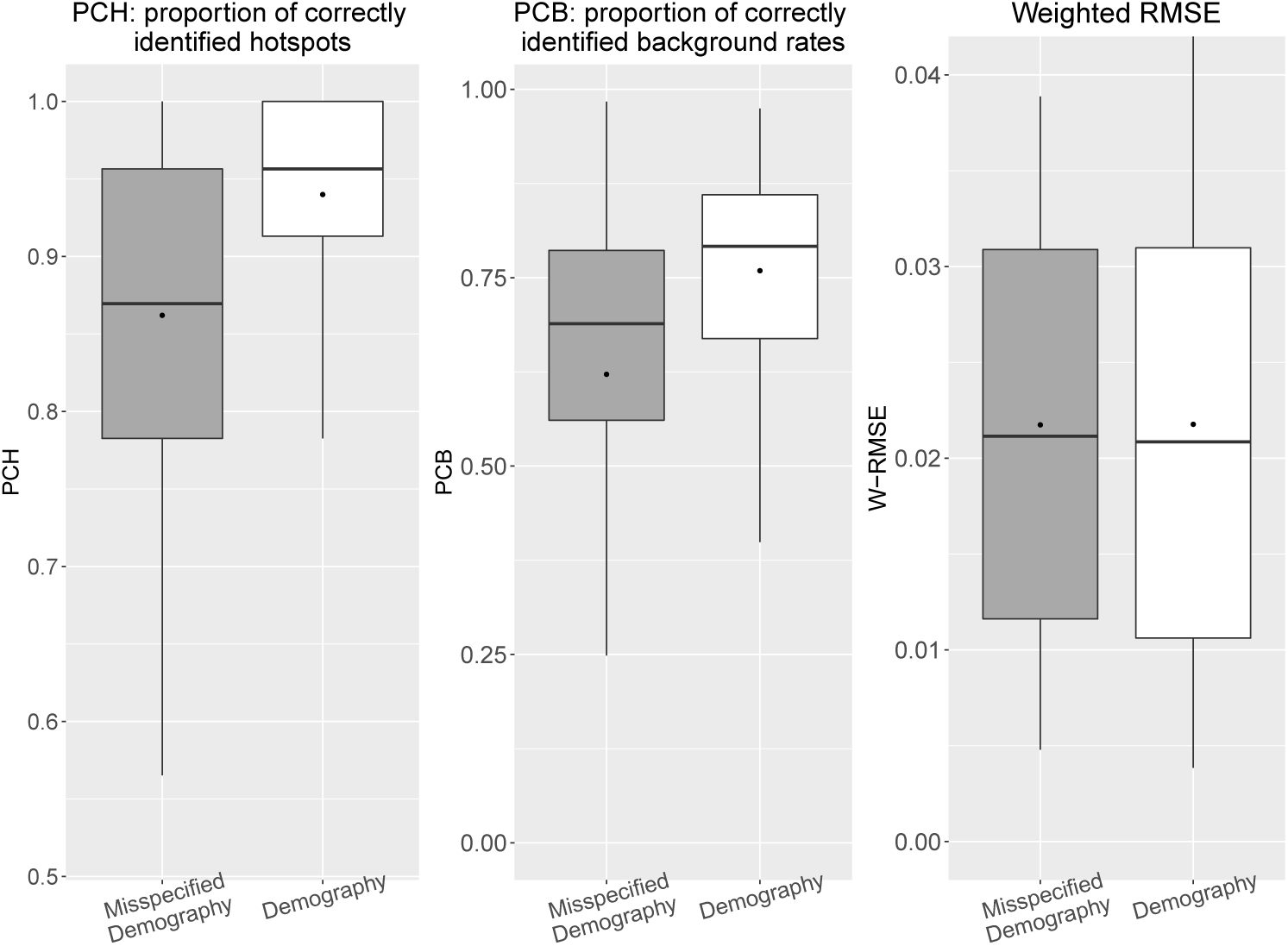
Performance of *LDJump* under the demographic model (2) (grey-boxes) compared with the results under *misspecified demography* (white-boxes), where a neutral model was incorrectly assumed. We set the segment lengths to 1kb for these comparisons and use the quantile of 0.35 in the bias correction. We provide box plots for the quality measures WAP (left), PCB (middle), and WMRSE (right).

### 3.5 Runtime

Obtaining estimates of recombination can be computationally demanding, especially for a larger number of sequences, and separate analyses for several populations. Hence, we also provide a comparison with respect to runtime (in seconds) between *LDhat, LDhat2, LDhelmet, FastEPRR*, and *LDJump*. We first consider simple setups using our simulated sequences of length 20kb. Again, we looked at different block penalty choices, as well as at different numbers of atomic segments *k* for *LDJump* in Table 3. As summaries, we computed the mean (top), median (middle), and SD (bottom) of our measured runtimes. We can see that especially *LDhat2* and *LDhelmet* run ten to fifty times longer than *LDJump* and *FastEPRR*. While being only slightly slower, we have seen before that *LDhat* leads to considerably less accurate estimates. *LDJump* turns out to be also faster than *FastEPRR* when the number of segments *k* is at least 15.

**Table 3:** Mean 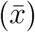, median (*x*_0.5_), and SD of runtime (in seconds) for *LDhat, LDhat2, LDhelmet*, *FastEPRR*, and *LDJump* under simple setups of length 20kb. For each method, separate columns provide values depending on either the block penalty for *LDhat, LDhat2, LDhelmet* (columns 2-4, 5-7, 8-10, respectively), the segment length (seg. length) for *FastEPRR* (columns 11-14) or the number of predefined segments *k* on which *LDJump* was applied (columns 15-18).

|  | *LDhat* (*bpen*) |  |  | *LDhat2* (*bpen*) |  |  | *LDhelmet* (*bpen*) |  |  | *FastEP RR* (seg. length) |  |  |  | *LDJump* (*k*) |  |  |  |
| --- | --- | --- | --- | --- | --- | --- | --- | --- | --- | --- | --- | --- | --- | --- | --- | --- | --- |
|  | 0 | 5 | 50 | 0 | 5 | 50 | 0 | 5 | 50 | 500 | 1000 | 1500 | 2000 | 10 | 20 | 25 | 30 |
| $\bar{x}$ | 35 | 56 | 156 | 751 | 3333 | 3260 | 1281 | 1368 | 1958 | 139 | 113 | 85 | 65 | 94 | 59 | 53 | 50 |
| $x_{0.5}$ | 34 | 55 | 138 | 735 | 3315 | 3261 | 849 | 936 | 1575 | 121 | 131 | 98 | 77 | 95 | 59 | 53 | 50 |
| SD | 6 | 7 | 70 | 273 | 999 | 977 | 1034 | 1042 | 1125 | 84 | 40 | 28 | 21 | 25 | 13 | 11 | 9 |

In Table 4 we show the mean, median, and SD of runtimes in seconds based on *natural* setups. On average *LDJump* turns out to be about ten to twenty times faster than *LDhat2*. Choosing larger values of *k* reduces the runtime for *LDJump*. Due to the faster computation of certain summary statistics, runtime was reduced by about 50% when going from 500 to 2000 segments.

**Table 4:**
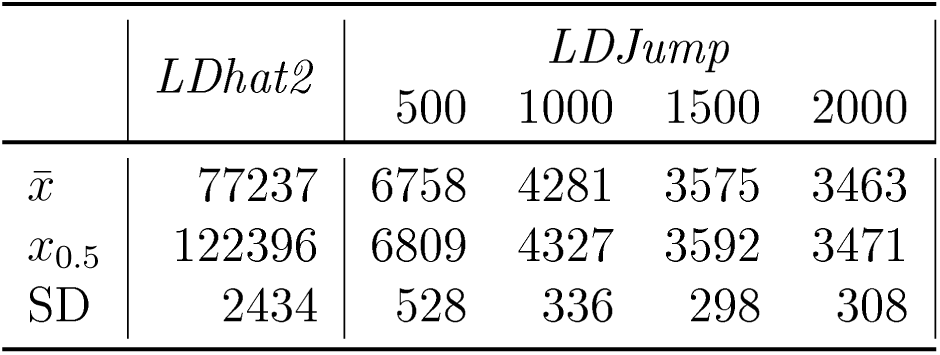
Mean 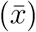, median (*x*_0.5_), and SD of the runtimes (in seconds) for *LDhat2* and *LDJump* under our natural setups. For *LDJump* we provide values depending on the number of predefined segments.

|  | *LDhat2* | *LDJump* |  |  |  |
| --- | --- | --- | --- | --- | --- |
|  |  | 500 | 1000 | 1500 | 2000 |
| $\bar{x}$ | 77237 | 6758 | 4281 | 3575 | 3463 |
| $x_{0.5}$ | 122396 | 6809 | 4327 | 3592 | 3471 |
| SD | 2434 | 528 | 336 | 298 | 308 |

In contrast to our approach. It turns out that the runtimes strongly depend on the underlying recombination rates with *LDhat2*, leading to a considerable difference between the median and mean of times. In supplementary material (section 4), we compare the runtimes for various background rates and different values of *k*. Overall, *LDJump* provides a particularly attractive combination of performance and runtime.

### 3.6 Validation of *LDJump* Computed Hotspots with Active Recombination Hotspots

We first tested our algorithm on a 103kb region on human chromosome 21. Therefore, we sampled the region between SNPs rs10622653 and rs2299784 containing the PCP4 gene, because this is the only region in which recombination has been characterized at high resolution by sperm typing for such a long continuous stretch [Tiemann-Boege et al., 2006]. Taking data from [The 1000 Genomes Project Consortium, 2015], we randomly chose 50 individuals for each of 4 subpopulations from 4 European regions (TSI, FIN, IBS, GBR). The data has been reformatted from `vcf`-format to `fasta`-files with the `R` packages [Knaus and Grünwald, 2017, Paradis et al., 2004] using two sequences per (diploid) sample and the reference sequence 80.37 (GRCH37) from [The 1000 Genomes Project Consortium, 2015]. We applied *LDJump* with a segment length of 1kb and chose the 35%-quantile for the bias-correction.

In the region from 60-100kb, the estimated recombination maps across populations (using a lookup table of 100 sequences and *θ* of 0.005) coincide well to the map obtained experimentally using sperm typing in [Tiemann-Boege et al., 2006] (see panel A of Figure 9). However, we also find population specific differences in the detected hotspots. We then compared this region with the double strand break (DSB) maps (representing active recombination hotspots) from [Pratto et al., 2014], see panel C of Figure 9. The hotspots inferred by *LDJump* in this region (60-100kb) also agree with the DSB activity. However, *LDJump* additionally estimated hotspots before the PCP4 gene around position 45kb. These hotspots were also found by other LD-based algorithms [McVean et al., 2004, Li and Stephens, 2003], see panel B of Figure 9 [Tiemann-Boege et al., 2006].

**Figure 9:**
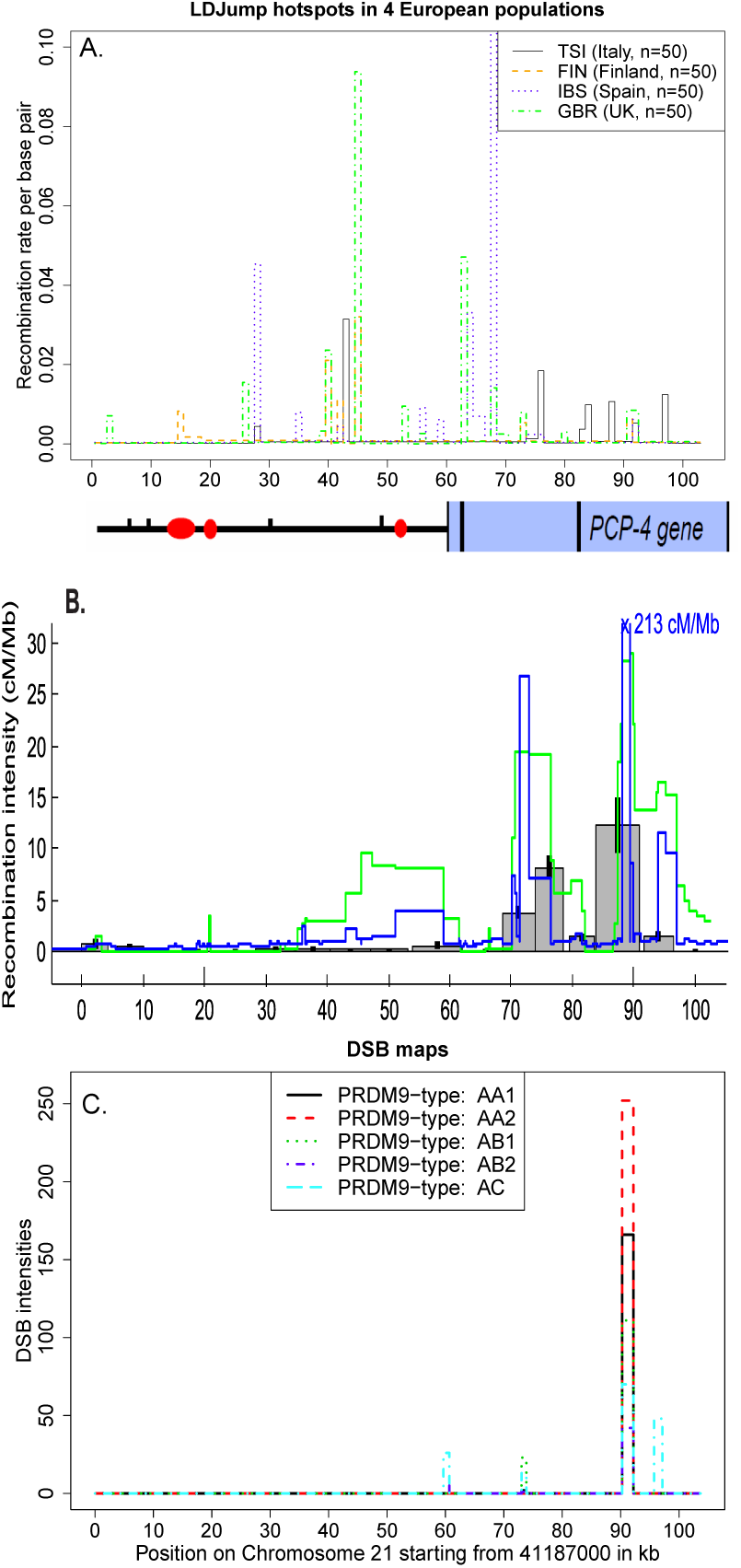
A: Estimated recombination map of 4 different European populations (Italy, Finland, Spain, and United Kingdom) on Chromosome 21:41187000-41290679 (GRCH37), including the PCP4-Gene. B: Estimated recombination map of the same 103kb region including PCP4 on Chromosome 21 taken from [Tiemann-Boege et al., 2006] based on sperm typing 13 intervals ~ 5kb in size (grey boxes). Moreover, the estimated recombination maps of *LDhat* (green) and Hotspotter (blue) are provided. C: Recombination maps based on measured double strand break (DSB) intensities for five different individuals representing active recombination from [Pratto et al., 2014].

We further tested the performance of *LDJump* within a larger genomic region to validate our method. For this purpose, we applied *LDJump* to the entire chromosome 16, and consider separate samples of 50 sequences from four populations (GBR, TSI, IBS, FIN) taken from [The 1000 Genomes Project Consortium, 2015]. For the data preparation we used the software package *vcftools* [Danecek et al., 2011] and then ran a parallel version of *LDJump* with segment lengths of 1kb for each population recombination map. We obtained these results in less than 2 days using in total 15 cores of an Intel Xeon E5-2630v3 2.4 1866, with 64GB DDR4-2133 RAM.

In panel A of Figure 10 we show the estimated recombination maps under the demography model (2) for chromosome 16 with the Italian population (TSI) in black, the Finnish sample in dashed red (FIN), the Spanish sample (IBS) in dotted green, and the British population (GBR) in dash-dotted blue. When we ignore demography and applied *LDJump* under a neutral scenario, we obtained hotspots with unrealistically high intensities (up to values of 70). Demography model (2) is rather simple, and we stress that *LDJump* could also be applied under any demographic scenario by training the regression model with a suitable setup. Overall, we observe population specific hotspots, but also hotspots present in more than one population as is also observed in genome-wide DSB maps (Figure 10, panel B) [Pratto et al., 2014].

**Figure 10:**
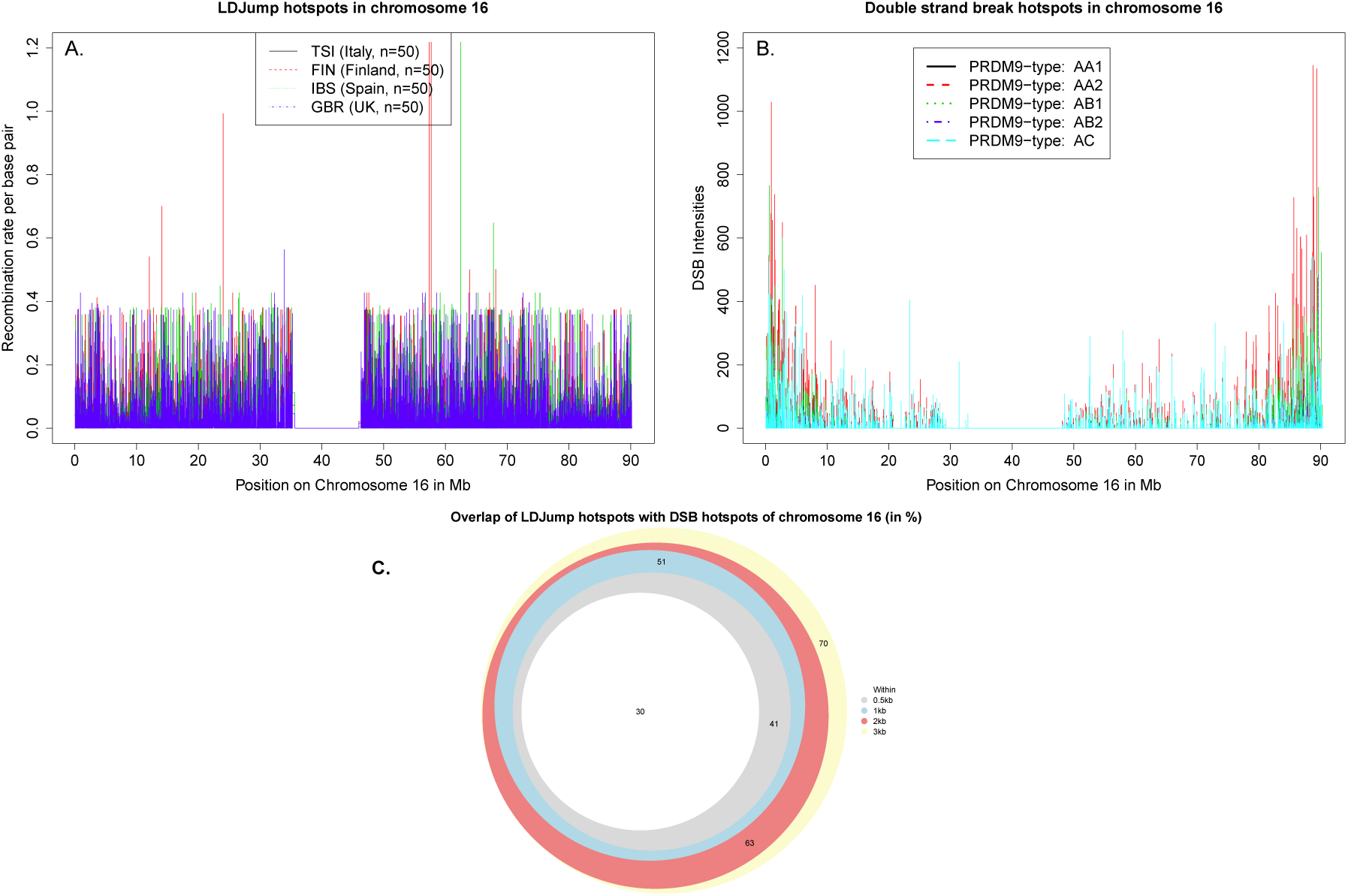
A: Estimated recombination map for chromosome 16 of four European populations with 50 randomly sampled individuals of the 1000 Genomes Project using *LDJump* under the demography model and segment lengths of 1kb. The results of the Italian sample are plotted in black, the Finnish sample in dashed red, the Spanish population in dotted green and the British one in dash-dotted blue. B: Double-strand break maps taken from [Pratto et al., 2014] of chromosome 16 for five individuals with different PR,DM9-types. Here, the different colors and line types represent different individuals (solid-black: AA1, dashed-red: AA2, dotted-green: AB1, dash-dotted-blue: AB2, long-dashed-cyan: AC). C: Overlap between detected DSB hotspots and the hotspots identified by *LDJump*. With *LDJump*, we define hotspots as regions with more than five times the estimated background rate. The DSB hotspots were taken from [Pratto et al., 2014]. We looked at overlaps between *LDJump* and the DSB hotspots that occurred with at least one European population (white areas). To assess the level of accuracy, we added segments of length 0.5 (grey), 1 (cyan), 2 (red), and 3 (yellow) kb left and right to the DSB region boundaries. The comparison is performed for all PRDM9-types considered in [Pratto et al., 2014]. The total number of DSB hotspots of all PRDM9-types is 2889 of which 866 were not detected by *LDJump*.

Furthermore, we evaluated the agreement of the estimated recombination hotspot locations using *LDJump* with the DSB map hotspots. For identifying *LDJump* hotspots we use a simple heuristic to define the average background rate. More specifically, we chose the mean of all estimates 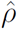 that fall below the median. This should give a downward biased estimate. With *LDJump*, we againd defined regions with more than five-fold the estimated background rate as hotspots. The DSB hotspots were selected by making use of the indicator variables provided by [Pratto et al., 2014]. Given that DSB hotspots are very narrow, yet the resolution of DSB into a crossover can occur with 3-5 kb, we added segments of different length (0, 0.5, 1, 2, 3 kb) left and right to the DSB hotspot regions and calculated the respective number of detected hotspots per PRDM9-type. The total number of DSB hotspots for AA1, AA2, AB1, AB2, and AC is 2889. We counted a hotspot as jointly detected, if an overlap between DSB hotspot and a *LDJump* hotspot occurred in at least one of the four populations (FIN, IBS, GBR, TSI). We display the number of jointly detected hotspots (augmented by segments of different lengths) via a Venn diagram in panel C of Figure 10. Notice that the number of hotspots estimated by *LDJump* for all considered populations is in total 31423, and therefore approximately 10 fold higher than the number of DSB-hotspots. Our analysis shows that on average about 70% of the DSB-hotspots (when adding 3kb segments to these regions) overlapped with at least one of the estimated *LDJump* population hotspots. These proportions go in line with the comparison of LD-based recombination maps and DSB hotspots in [Pratto et al., 2014] with an overlap of 56%.

## 4 Discussion

We introduced a new method called *LDJump* to estimate heterogeneous recombination rates along chromosomes from population genetic data. Our approach splits a given DNA sequence into segments of proper length in a first step. Subsequently, we use a generalized additive regression model to estimate the constant recombination rates per segment. Then, we apply a simultaneous multiscale change-point estimator (SMUCE) to estimate the breakpoints in the recombination rates across the sequence. We provide detailed comparisons of our method with the recent reversible jump MCMC methods *LDhat(2)* and *LDhelmet* as well as the regression based method *FastEPRR*. Our estimates are very fast, perform favourably in the detection of hotspots, and show similar accuracy levels as the best available competitor for *simple* and *natural* setups, respectively. These comparisons show that *LDJump* is a powerful tool to explore recombination rates in organisms with narrow recombination hotspots; for example, PRDM9 defined hotspots in most mammals (reviewed in [Tiemann-Boege et al., 2017]).

We validated our method by computing hotspots in several human populations and compared the estimated hotspots with recombination intensities measured by sperm-typing and double strand break maps. These computations revealed population specific hotspots in the region surrounding the PCP4-gene located on chromosome 21. Although on a small scale (103b) *LDJump* computed hotspots mainly agree with hotspots detected at high resolution with sperm typing and Chip immuno-precipitation (DSB map); we also observed a region with little congruence at position ~45kb. Given the lack of active recombination at position ~45kb (absence of this hotspot in sperm typing and in the DSB maps for the 2 European donors carrying the PRDM9 allele A, as well as the donor with African descent (carrying the PRDM9 allele C)), we hypothesize that this estimated hotspot might represent a historical hotspot that got extinct. Alternatively it could be a population-specific hotspot given that its intensity varies among different European populations. In order to test this latter hypothesis, active recombination maps from different populations would be needed. Experimental data (e.g. hotspot at position 95kb present only in the individual with a PRDM9 C allele) suggest that the intensity of hotspots might vary also within populations.

Differences between hotspot rates estimated from LD patterns compared to estimates based on sperm typing have also been observed by [Jeffreys and Neumann, 2009]. This might be caused by the short life-span of hotspots and their rapid evolution in intensity and genomic position among populations and species [Coop and Myers, 2007, Myers et al., 2010, Jeffreys et al., 2013]. In fact, only ~56% of historical hotspots determined by LD agree with genome-wide DSB maps [Pratto et al., 2014]. Our large-scale validation on chromosome 16 shows that about 70% of the DSB-hotspots were also found by *LDJump* using four European populations. Fine-scale population specific differences with respect to recombination events have also been highlighted in studies such as [Kong et al., 2010, Berg et al., 2011, Fledel-Alon et al., 2011, Pratto et al., 2014]. Given all this, our observed differences are likely due to underlying biological features. We have implemented our approach as an `R`-package called *LDJump*, which can be freely downloaded from https://github.com/PhHermann/LDJump. In our simulations, we obtained particularly good results when applying our method with segment lengths of 1kb and a bias correction using the default quantile of 0.35.

In conclusion, *LDJump* is a fast algorithm which is able to detect narrow hotspots at high accuracy using segments of approximately 1kb length. Moreover, we also show that *LDJump* can be applied on populations under demography. We validated our method on a 103kb region of human chromosome 21 as well as the whole chromosome 16 and found a good congruence by comparing *LDJump* hotspots with recombination hotspots measured with sperm typing or Chip immuno-precipitation (DSB map).

## Acknowledgements

We are grateful to Kerstin Gärtner and Renato Pereira Salazar for their assistance with the software packages *LDhat* and *LDhelmet* as well as the data acquisition for the application, respectively. We are thankful to Katharina Sallinger, Bettina Grün, Renato Pereira Salazar, and Theresa Schwarz for their helpful comments. This work was supported by the ‘Austrian Science Fund’ (FWF) P27698-B22 to I.T-B., and the DOC Fellowship of the Austrian Academy of Sciences (24529) to A.H.

## Data Accessibility Statement

*LDJump* has been implemented as an R package which can be downloaded and installed from Github (https://github.com/PhHermann/LDJump). We also provide example files and a manual in this repository. We downloaded the data of chromosome 16 from ftp://ftp.1000genomes.ebi.ac.uk/vol1/ftp/release/20130502 and uploaded an R-script with details on the data management as well as the hotspot locations for the estimated population recombination maps to Github (https://github.com/PhHermann/Hermann_et_al_2018_LDJump). We download the data for the application on chromosome 21 from http://phase3browser.l000genomes.org/Homo_sapiens/Location/0verview?r=21:41187000-41290679 using the first 50 samples of the four European populations IBS, GBR, TSI, and FIN.

We provide details on the regression model, bias correction, choice of segment lengths, detailed quality assessments and runtime comparisons in the *supplementary material*.

## Author Contributions

PH and AF designed the model and implemented the model into the R package. PH and AF focused on the statistical aspects and ITB and AH on the biological aspects. All authors wrote and commented on the manuscript.

## References

Arbeithuber, B., Betancourt, A. J., Ebner, T., and Tiemann-Boege, I. (2015). Crossovers are associated with mutation and biased gene conversion at recombination hotspots. Proceedings of the National Academy of Sciences, 112(7):2109–2114.

Arenas, M., Lopes, J. S., Beaumont, M. A., and Posada, D. (2015). CodABC: A Computational Framework to Coestimate Recombination, Substitution, and Molecular Adaptation Rates by Approximate Bayesian Computation. Molecular Biology and Evolution, 32(4):1109–1112.

Arnheim, N., Calabrese, P., and Tiemann-Boege, I. (2007). Mammalian Meiotic Recombination Hot Spots. Annual Review of Genetics, 41(1):369–399.

Auton, A., Fledel-Alon, A., Pfeifer, S., Venn, O., Segurel, L., Street, T., Leffler, E. M., Bowden, R., Aneas, I., Broxholme, J., Humburg, P., Iqbal, Z., Lunter, G., Maller, J., Hernandez, R. D., Melton, C., Venkat, A., Nobrega, M. A., Bontrop, R., Myers, S., Donnelly, P., Przeworski, M., and McVean, G. (2012). A Fine-Scale Chimpanzee Genetic Map from Population Sequencing. Science, 336(6078): 193–198.

Auton, A. and McVean, G. (2007). Recombination rate estimation in the presence of hotspots. Genome Research, 17(8): 1219–1227.

Batorsky, R., Kearney, M. F., Palmer, S. E., Maldarelli, F., Rouzine, I. M., and Coffin, J. M. (2011). Estimate of effective recombination rate and average selection coefficient for HIV in chronic infection. Proceedings of the National Academy of Sciences of the United States of America, 108(14):5661–6.

Baudat, F., Imai, Y., and de Massy, B. (2013). Meiotic recombination in mammals: localization and regulation. Nature reviews. Genetics, 14(11):794–806.

Berg, I. L., Neumann, R., Lam, K.-W. G., Sarbajna, S., Odenthal-Hesse, L., May, C. A., and Jeffreys, A. J. (2010). PRDM9 variation strongly influences recombination hot-spot activity and meiotic instability in humans. Nature Genetics, 42(10):859–863.

Berg, I. L., Neumann, R., Sarbajna, S., Odenthal-Hesse, L., Butler, N. J., and Jeffreys, A. J. (2011). Variants of the protein PRDM9 differentially regulate a set of human meiotic recombination hotspots highly active in African populations. Proc Natl Acad Sci U S A, 108 (30): 12378–12383.

Box, G. E. P., and Cox, D. R. (1964). An analysis of transformations. Journal of the Royal Statistical Society. Series B (Methodological, pages 211–252.

Chan, A. H., Jenkins, P. A., and Song, Y. S. (2012). Genome-Wide Fine-Scale Recombination Rate Variation in Drosophila melanogaster. PLoS Genetics, 8(12):e1003090.

Coop, G. and Myers, S. R. (2007). Live Hot, Die Young: Transmission Distortion in Recombination Hotspots. PLoS Genetics, 3(3):e35.

Coop, G., Wen, X., Ober, C., Pritchard, J. Κ., and Przeworski, M. (2008). High-Resolution Mapping of Crossovers Reveals Extensive Variation in Fine-Scale Recombination Patterns Among Humans. Science, 319(5868): 1395–1398.

Danecek, P., Auton, A., Abecasis, G., Albers, C. A., Banks, E., DePristo, M. A., Handsaker, R. E., Lunter, G., Marth, G. T., Sherry, S. T., McVean, G., and Durbin, R. (2011). The variant call format and VCFtools. Bioinformatics, 27(15):2156–2158.

Fearnhead, P. and Donnelly, P. (2001). Estimating recombination rates from population genetic data. Genetics, 159: 1299–1318.

Fearnhead, P. and Donnelly, P. (2002). Approximate Likelihood Methods for Estimating Local Recombination Rates. Journal of the Royal Statistical Society Series B (Statistical Methodology), 64(4):657–680.

Fledel-Alon, A., Leffler, E. M., Guan, Y., Stephens, M., Coop, G., and Przeworski, M. (2011). Variation in human recombination rates and its genetic determinants. PLoS ONE, 6(6).

Frick, K., Munk, A., and Sieling, H. (2014). Multiscale change-point inference. Journal of the Royal Statistical Society: Series B, 76(3):495–580.

Futschik, A., Hotz, T., Munk, A., and Sieling, H. (2014). Multiscale DNA partitioning: Statistical evidence for segments. Bioinformatics, 30(16):2255–2262.

Gao, F., Ming, C., Hu, W., and Li, H. (2016). New Software for the Fast Estimation of Population Recombination Rates (FastEPRR) in the Genomic Era. G3 (Bethesda, Md.), 6(6): 1563–1571.

Gärtner, K. and Futschik, A. (2016). Improved Versions of Common Estimators of the Recombination Rate. Journal of Computational Biology, 23(9):756–768.

Halldorsson, B. V., Hardarson, M. T., Kehr, B., Styrkarsdottir, U., Gylfason, A., Thorleifsson, G., Zink, F., Jonasdottir, A., Jonasdottir, A., Sulem, P., Masson, G., Thorsteinsdottir, U., Helgason, A., Kong, A., Gudbjartsson, D. F., and Stefansson, K. (2016). The rate of meiotic gene conversion varies by sex and age. Nat Genet, 48(11):1377–1384.

Haubold, B. and Pfaffelhuber, P. (2013). ms2dna, v. 1.16: Convert Simulated Haplotype Data to DNA Sequences.

Hill, W. G., and Robertson, A. (1968). Linkage disequilibrium in finite populations. Theoretical and Applied Genetics, 38(6):226–231.

Hotz, T. and Sieling, H. (2016). stepR: Fitting Step-Functions.

Hudson, R. R. (1987). Estimating the recombination parameter of a finite population model without selection. Genetical research, 50(2007):245–250.

Hudson, R. R. (2001). Two-locus sampling distributions and their application. Genetics, 159(4):1805–1817.

Hudson, R. R., and Kaplan, N. L. (1985). Statistical properties of the number of recombination events in the history of a sample of DNA sequences. Genetics, 111(1): 147–164.

Jeffreys, A. J., Cotton, V. E., Neumann, R., and Lam, K.-W. G. (2013). Recombination regulator PRDM9 influences the instability of its own coding sequence in humans. Proceedings of the National Academy of Sciences, 110(2):600–605.

Jeffreys, A. J., and Neumann, R. (2009). The rise and fall of a human recombination hot spot. Nature genetics, 41(5):625–9.

Johnston, H. R., and Cutler, D. J. (2012). Population demographic history can cause the appearance of recombination hotspots.

Jombart, T. (2008). adegenet: a R package for the multivariate analysis of genetic markers. Bioinformatics, 24(11): 1403–1405.

Kamm, J. A., Spence, J. P., Chan, J., and Song, Y. S. (2016). Two-Locus Likelihoods Under Variable Population Size and Fine-Scale Recombination Rate Estimation. Genetics, 203(3):1381 LP–1399.

Kauppi, L., Jeffreys, A. J., and Keeney, S. (2004). Where the crossovers are: recombination distributions in mammals. Nature Reviews Genetics, 5(6):413–424.

Knaus, B. J., and Grünwald, N. J. (2017). vcfr: a package to manipulate and visualize variant call format data in R. Molecular Ecology Resources, 17(1):44–53.

Kong, A., Thorleifsson, G., Gudbjartsson, D. F., Masson, G., Sigurdsson, A., Jonasdottir, A., Walters, G. B., Jonasdottir, A., Gylfason, A., Kristinsson, Κ. T., Gudjonsson, S. A., Frigge, M. L., Helgason, A., Thorsteinsdottir, U., and Stefansson, K. (2010). Fine-scale recombination rate differences between sexes, populations and individuals. Nature, 467(7319): 1099–1103.

Kuhner, M. Κ. (2006). LAMARC 2.0: maximum likelihood and Bayesian estimation of population parameters. Bioinformatics, 22(6):768–770.

Kuhner, M. K., Yamato, J., and Felsenstein, J. (2000). Maximum likelihood estimation of recombination rates from population data. Genetics, 156(3):1393–1401.

Li, N. and Stephens, M. (2003). Modeling Linkage Disequilibrium and Identifying Recombination Hotspots Using Single-Nucleotide Polymorphism Data. Genetics, 165(4):2213–2233.

Lin, Κ., Futschik, A., and Li, H. (2013). A Fast Estimate for the Population Recombination Rate Based on Regression. Genetics, 194(2):473–484.

Martin, D. P., Murrell, B., Golden, M., Khoosal, A., and Muhire, B. (2015). RDP4: Detection and analysis of recombination patterns in virus genomes. Virus Evolution, 1(1).

McVean, G. A. T., Awadalla, P., and Fearnhead, P. (2002). A coalescent-based method for detecting and estimating recombination from gene sequences. Genetics, 160(3):1231–41.

McVean, G. A. T., Myers, S. R., Hunt, S., Deloukas, P., Bentley, D. R., and Donnelly, P. (2004). The fine-scale structure of recombination rate variation in the human genome. Science, 304(5670):581–584.

Myers, S., Bottolo, L., Freeman, C., McVean, G., and Donnelly, P. (2005). A Fine-Scale Map of Recombination Rates and Hotspots Across the Human Genome. Science, 310(5746):321–321.

Myers, S., Bowden, R., Tumian, A., Bontrop, R. E., Freeman, C., MacFie, T. S., McVean, G., and Donnelly, P. (2010). Drive Against Hotspot Motifs in Primates Implicates the PRDM9 Gene in Meiotic Recombination. Science, 327(5967):876–879.

Myers, S., Freeman, C., Auton, A., Donnelly, P., and McVean, G. (2008). A common sequence motif associated with recombination hot spots and genome instability in humans. Nature Genetics, 40(9): 1124–1129.

Myers, S. R., and Griffiths, R. C. (2003). Bounds on the minimum number of recombination events in a sample history. Genetics, 163(1):375–394.

Niu, Y. S., Hao, N., and Zhang, H. (2016). Multiple Change-Point Detection: A Selective Overview. Statistical Science, 31(4):611–623.

Paradis, E., Claude, J., and Strimmer, K. (2004). A[PE]: analyses of phylogenetics and evolution in [R] language. Bioinformatics, 20: 289–290.

Pérez-Losada, M., Arenas, M., Galán, J. C., Palero, F., and González-Candelas, F. (2015). Recombination in viruses: Mechanisms, methods of study, and evolutionary consequences. Infection, Genetics and Evolution, 30: 296–307.

Pratto, F., Brick, K., Khil, P., Smagulova, F., Petukhova, G. V., and Camerini-Otero, R. D. (2014). Recombination initiation maps of individual human genomes. Science, 346(6211).

R Development Core Team (2017). R: A language and environment for statistical computing.

Reid, N. (2013). Aspects of likelihood inference. Bernoulli, 19(4):1404–1418.

Sabeti, P. C., Schaffner, S. F., Fry, B., Lohmueller, J., Varilly, P., Shamovsky, O., Palma, A., Mikkelsen, T. S., Altshuler, D., and Lander, E. S. (2006). Positive natural selection in the human lineage. Science (New York, N.Y.), 312(5780): 1614–20.

Smagulova, F., Gregoretti, I. V., Brick, Κ., Khil, P., Camerini-Otero, R. D., and Petukhova, G. V. (2011). Genome-wide analysis reveals novel molecular features of mouse recombination hotspots. Nature, 472(7343):375–378.

Smith, N. G. C. (2005). A Comparison of Three Estimators of the Population-Scaled Recombination Rate: Accuracy and Robustness. Genetics, 171(4):2051–2062.

Staab, P. R., Zhu, S., Metzler, D., and Lunter, G. (2014). Scrm: Efficiently simulating long sequences using the approximated coalescent with recombination. Bioinformatics, 31(10):1680–1682.

Tajima, F. (1989). Statistical method for testing the neutral mutation hypothesis by DNA polymorphism. Genetics, 123(3):585–595.

The 1000 Genomes Project Consortium (2015). A global reference for human genetic variation. Nature, 526(7571):68–71.

Tiemann-Boege, I., Calabrese, P., Cochran, D. M., Sokol, R., and Arnheim, N. (2006). High-Resolution Recombination Patterns in a Region of Human Chromosome 21 Measured by Sperm Typing. PLoS Genetics, 2(5):e70.

Tiemann-Boege, I., Schwarz, T., Striedner, Y., and Heissl, A. (2017). The consequences of sequence erosion in the evolution of recombination hotspots. Philosophical Transactions of the Royal Society B: Biological Sciences, 372(1736):20160462.

Varin, C., Reid, N., and Firth, D. (2011). An Overview of Composite Likelihood Methods. Statistica Sinica, 21(1):5–42.

Wall, J. D. (2000). A Comparison of Estimators of the Population Recombination Rate. Mol. Biol. Evol, 17(1): 156–163.

Wall, J. D. (2004). Estimating recombination rates using three-site likelihoods. Genetics, 167(3):1461–1473.

Warnes, G., Gorjanc, G., Leisch, F., and Man, M. (2013). genetics: Population Genetics.

Wilson, D. J., and McVean, G. (2006). Estimating Diversifying Selection and Functional Constraint in the Presence of Recombination. Genetics, 172(3):1411–1425.

Wiuf, C. (2002). On the minimum number of topologies explaining a sample of DNA sequences. Theoretical Population Biology, 62(4):357–363.

Wood, S. N. (2011). Fast stable restricted maximum likelihood and marginal likelihood estimation of semiparametric generalized linear models. Journal of the Royal Statistical Society: Series B (Statistical Methodology), 73(1):3–36.

